# Energy Landscape of the Domain Movement in *Staphylococcus aureus* UDP-N-acetylglucosamine 2-epimerase

**DOI:** 10.1101/535138

**Authors:** Erika Chang de Azevedo, Alessandro S. Nascimento

## Abstract

*Staphylococcus aureus* is an important cause of resistant healthcare-associated infections. It has been shown that the Wall Teichoic Acid (WTA) may be an important drug target acting on antibiotic-resistant cells. The UDP-N-Acetylglucosamine 2-epimerase, MnaA, is one of the first enzymes on the pathway for the biosynthesis of the WTA. Here, detailed molecular dynamics simulations of *S. aureus* MnaA were used to characterize the conformational changes that occur in the presence of UDP and UDP-GlcNac and also the energetic landscape associated with these changes. Using different simulation techniques, such as ABMD and GAMD, it was possible to assess the energetic profile for the protein with and without ligands in its active site. We found that there is a dynamic energy landscape that has its minimum changed by the presence of the ligands, with a closed structure of the enzyme being more frequently observed for the bound state while the unbound enzyme favors an opened conformation. Further structural analysis indicated that positively charged amino acids associated with UDP and UDP-GlcNac interactions play a major role in the enzyme opening movement. Finally, the energy landscape profiled in this work provides important conclusions for the design of inhibitor candidates targeting *S. aureus* MnaA.

## Introduction

*Staphylococcus aureus* is a naturally occurring Gram-positive bacteria that can be often found in the skin and mucosa of humans and other animals (McMillan et al., 2016). Although usually harmless, this organism can cause a wide range of infections in humans, such as folliculitis, furuncles, and carbuncles, impetigo, mastitis, wound infections, and staphylococcal scalded skin syndrome. More dangerous pathological conditions caused by *S. aureus* include bacteremia, pneumonia, endocarditis, bone, and joint infections, and toxic shock syndrome (Peacock and Paterson, 2015). *S. aureus* is also an important cause of hospital infections. According to data collected by the CDC between 2011 and 2014, this microorganism was responsible for approximately 10% of the overall healthcare-associated infections analyzed in the United States (L.M. et al., 2016).

The β–lactams, such as methicillin and penicillin, are among the antibiotics considered as the first choice for the treatment of this Gram-positive bacterial infection. These antibiotics act on the peptidoglycan layer of the cell, inhibiting transglycosylases and transpeptidases, collectively known as penicillin-binding proteins or PBPs, that are responsible for the synthesis and crosslink of specific polymers of the cell wall (Farha et al., 2013). However, the treatment of Staphylococcal infections is becoming more challenging due to the emergence of drug-resistant strains (Lowy, 1998). These resistant strains were first reported only a few years after the introduction of penicillin and have increased in number of occurrences faster after the introduction of new antibiotics (Sewell and Brown, 2014). The CDC data, taken from 4,515 reporting hospitals and over 365,000 healthcare-associated infections, show that about 50% of the infections caused by *S. aureus* in healthcare units had isolated strains that were reported to be resistant to the β-lactam antibiotics like oxacillin, methicillin, and cofoxitin (L.M. et al., 2016).

*S. aureus* antibiotic resistance has a direct impact in the human lives but also accounts for an important burden in the health systems. Data taken in 2007 from 31 countries that participate in the European Antimicrobial Resistance Surveillance System (EARSS) showed that 27,711 bloodstream infections caused by MRSA resulted in 5,503 excess deaths and 255,683 excess hospital days, thus generating in a burden of 44 million Euros (63.1 million dollars) (de Kraker et al., 2011).

The emergence of antibiotic resistance highlights the need for new drugs or even new ways to re-sensitize resistant bacterium. In this context, the cell wall remains as an interesting target for the discovery of new drugs and agents that fight β–lactam resistance (Farha et al., 2013). The cell wall of Gram-positive bacteria, such as *S. aureus*, is typically composed by peptidoglycan and wall teichoic acid (WTA). The WTA is located in the outer layer of the peptidoglycan and its role is associated the maintenance of the cell shape, virulence, coordination of the peptidoglycan synthesis, biofilm formation and, more recently, it has been linked to antibiotic resistance (Brown et al., 2012; Roemer et al., 2013; Sewell and Brown, 2014; Sobhanifar et al., 2016). In *S. aureus*, the WTA is typically made of a poly(ribitol-phosphate) polymer that is synthesized by the sequential action of several enzymes, called *Tar* (Teichoic acid ribitol) enzymes, starting with the glycosyltransferases TarA and TarO. Another important enzyme involved in the biosynthesis of WTA and in the *S. aureus* resistance to β-lactams is the N-acetylglucosamine 2-epimerase or MnaA (Mann et al., 2016). This enzyme interconverts UDP-N-acetylglucosamine and UDP-N-acetylmannosamine, which are the substrates for TarO and TarA, respectively, thus regulating the balance of the starting points of the Tar biosynthesis pathway (Mann et al., 2016).

The *S. aureus* MnaA (SaMnaA) binds simultaneously to a UDP molecule and to the substrate UDP-N-acetylglucosamine (UDP-GlcNac) or UDP-N-acetylmannosamine (UDP-ManNac) (Mann et al., 2016; Velloso et al., 2008). Analysis of *S. aureus* MnaA (SaMnaA) crystal structure (Mann et al., 2016) and comparison with previously determined crystal structures revealed that the protein exhibits a domain movement upon binding the substrate, defining at least three observable conformations for the enzyme as opened, semi-closed and closed (Velloso et al., 2008). The opened state was observed in the absence of the substrate or product in the active site. The semi-closed state is observed in the presence of UDP. The closed state is typically observed when the protein has both UDP and UDP-GlcNac bound in its active site (Mann et al., 2016; Velloso et al., 2008). Interestingly, this movement between domains is also observed for several enzymes, including some enzymes of the WTA biosynthesis, such as TarM (Brown et al., 2012).

Here, we evaluated the domain movement of SaMnaA at the atomic level using different molecular dynamics simulations and assessing the free energy landscape associated to such movement. The simulations showed that the protein has a shift in the minimum of the free energy landscape profile when associated with the ligands UDP and UDP-GlcNac. Additionally, a major driving force that causes this shift seems to come from the electrostatic interactions between the enzyme and the substrate. These results can lead to a binding energy profile that indicates a dynamic energy landscape (Kumar et al., 2000) for the open/close movement of the protein, relevant for the enzymatic action.

## Materials and Methods

### Protein and Ligand Parameterization

The chain A of the *S. aureus* MnaA structure (PDB ID: 5ENZ) was parameterized using AMBER FF14SB force-field (Maier et al., 2015). Hydrogen atoms were added and the histidine protonation states determined at the optimum pH of 7.0 using the H++ server (Anandakrishnan et al., 2012; Bashford and Karplus, 1990). The protein was immersed in a water box using TIP3P water model (Price and Brooks, 2004) and sodium ions were also included to keep the systems electrically neutral. UDP and UDP-GlcNac atomic charges were calculated using the Hartree-Fock method, with 6-31G* basis set as available in Gaussian 09 (Rassolov et al., 2001). The obtained atomic charges were then refitted with RESP (Restrained Electrostatics Potential) (Comell et al., 1993) procedure using ANTECHAMBER (Wang et al., 2006). The Lennard-Jones parameters for these ligands defined using GAFF force field (Wang et al., 2004). The crystal structure of SaMnaA (PDB Id 5ENZ (Mann et al., 2016)) was used as the starting structure for the APO MD simulations. The UDP and UDP-GlcNac bound structure (HOLO) was modeled using the chain A of the crystal structure of the SaMnaA (PDB ID: 5ENZ) and the crystal structure of *Bacillus anthracis* MnaA (BaMnaA) bound with UDP and UDP-GlcNac (PDB ID: 3BEO (Velloso et al., 2008)), using the program Modeller (Šali and Blundell, 1993; Webb and Sali, 2016). The purpose of this initial modeling was to generate a ‘bound-state’ (closed) conformation of SaMnaA with UDP and UDP-GlcNac coordinates from BaMnaA. The final systems comprised 45,519 and 55,479 atoms for the APO and HOLO system, respectively.

### Equilibrium Simulations

The initial energy of the APO system was minimized in 1,000 cycles of conjugated gradient minimization. The system was then heated to 300 K in 50 ps, keeping the volume constant. The density of the system was afterward equilibrated to 1 g/cm^3^ by keeping constant pressure at 1 atm in 400 ps. During the minimization, heating and density equilibration, harmonic restraints were applied to all residues, with weight of 2 kcal.mol^-1^. Å^-2^. Finally, the system was equilibrated for 2 ns, without restraints, and the productive simulation was performed for 1 μs in an NPT ensemble and, afterward, extended up to 5 μs. Periodic boundary conditions were applied using a 12 Å cutoff radius for truncating short-range potentials. The simulation protocol for the HOLO system followed a similar procedure, with 4000 cycles of steepest descent energy minimization followed by 4000 steps of conjugated gradient minimization. The system was then heated to 300 k in 50 ps, with constant volume. The density was equilibrated until 1 g/cm^3^ in 400 ps keeping constant pressure at 1 atm. For minimization, heating and density equilibration, a harmonic positional restraint was applied to the entire protein and for the two ligands with a weight of 2 kcal.mol^-1^.Å^-2^. Finally, the system was equilibrated for 10 ns and the productive simulation was performed for 1 μs without restraints. All the simulations were performed using AMBER16 (Case et al., 2005; Salomon-Ferrer et al., 2013).

### Gaussian Accelerated Molecular Dynamics

In Gaussian Accelerated Molecular Dynamics (GAMD) (Miao et al., 2015), the trajectory is propagated in a non-negative Gaussian boost potential. This potential is applied to reduce the potential barriers and accelerate the state transitions, thus enhancing the conformational sampling. When the system potential *V* is lower than a threshold E, a harmonic boost potential Δ*V* is introduced to the total energy. The energy boost Δ*V* is chosen to result in the highest acceleration as possible but still allowing an accurate reweighting and free energy profile estimation. As shown in Equation 1, Δ*V* is defined as a harmonic potential with force constant k, which depends on the constant σ_0_ (Equations 2 and 3) (Miao et al., 2015). In Equations 2 and 3, V_max_ and V_min_ are the maximum and minimum system potential collected before the GAMD begins. σ_v_ and V_av_ are the standard deviation and average of the system potential energy, respectively. As previously observed, σ_0_ must be chosen carefully in a compromise to get the best possible acceleration but still allowing an accurate reweighting of the accelerated potentials. One advantage associated with GAMD and with Accelerated MD (aMD) is the lack of a pre-chosen *reaction coordinate* or a *collective variable* (CV). Also, as pointed out by Miao and coworkers, GAMD preserves the original potential surface, allowing the computation of a PMF with less noise as compared to aMD (Miao et al., 2015). Here, we applied the boost potential to the dihedral and the total potential of the system, in a dual-boost approach. For both potentials, the σ_0_ was set to 3 kcal/mol, based on preliminary simulations with varying values for σ_0_, where 3 kcal/mol provided the higher acceleration.

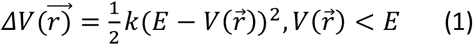

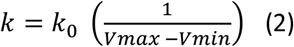

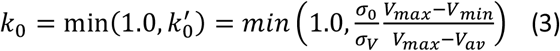

The resulting boost potential followed a Gaussian distribution that allowed a reweight and the calculation of a potential of mean force. The reweighting was made by obtaining the second-order of the cumulant expansion for the potential boost, as previously described (Miao et al., 2014). This calculation was made based on an opening angle computed between 10-80 degrees, with an angle increment of 4 degrees.

### Adaptively Biased Molecular Dynamics

The Adaptively Biased Molecular Dynamics (ABMD) is an umbrella sampling method similar, in spirit, to metadynamics. The biasing force compensates for the free energy gradient, reproducing the previously unknown free energy surface for the system analyzed in the direction of the collective variable (CV), chosen by the user (Babin et al., 2009). Here, we used the angle between the α-carbon of the residues S128, G349, and P213, describing the opening angle of the SaMnaA structure. The simulation was conducted with a well-tempered umbrella sampling method (Barducci et al., 2008), with pseudo-temperature T defined at 5,000 K. The total set of simulations used in this work are provided in Table 1.

**Table 1.** Simulations used in this work. The simulation methods and times used are shown.

| Method | Protein System | Time (ns) |
| --- | --- | --- |
| Equilibrium | APO | 1,000 + 4,000 |
| Equilibrium | HOLO | 1,000 |
| GAMD | APO | 3 x 500 |
| GAMD | HOLO | 3 x 250 |
| ABMD | APO | 3 x 250 |
| ABMD | HOLO | 3 x 250 |

### Structural Analysis

The APO system simulation of 5 μs was analyzed using the algorithms available in the pyEMMA library (Scherer et al., 2015). Only the carbon alpha coordinates from the protein were used in the analysis, and the lag time used in the time-lagged independent component analysis (TICA) was set 200 frames or 10 ns. The electrostatic forces between the C-terminal and N-terminal domains of the protein were conducted using the Delphi Force Web server (Li et al., 2017), where the first domain was defined as the protein region encompassing residues 1 to 181. Three representative states of the protein were used in the calculations, open, intermediate and closed, with the ligands considered as part of the C-terminal domain. Geometric analysis of the MD trajectories were carried out with CPPTRAJ (Roe and Cheatham, 2013) and interaction energies were computed with AmberEnergy++, a locally written program to compute interaction potential energies according to the AMBER force field (Nascimento, 2018).

## Results

The states defined as open, semi-closed and closed can be better characterized by the opening angle observed for protein structures available on the PDB. The UDP-GlcNac 2-epimerase of *Methanocaldococcus janaschii* structure was solved without ligands (PDB iD 4NEQ) (Chen et al., 2014) and with both UDP and UDP-GlcNac (PDB iD 4NES) (Chen et al., 2014). The first structure is in the opened state, with an opening angle (CV) of 54 degrees. Here, the opening angle is defined as the angle between the Cα atoms of the residues S128, G349, and P213 in SaMnaA, or residues in similar positions for the other epimerases. The second structure is in a closed state, with a CV angle of 28 degrees. The SaMnaA was in the semi-closed state, with CV angle of 31 degrees, for chain A. A description of the opening angles, defined by our chosen CV, for all the UDP-GlcNac 2-epimerases with available structures to date is given in Table 2.

From the analysis of the results shown in Table 2, it can be observed that (i) for crystal structures of UDP-N-acetylglucosamine 2-epimerases in the absence of ligands (1O6C, 1V4V, 3OT5, 4NEQ and 3DZC) the opening angle varies between 47 and 53°, indicating the tendency to a opened state without substrate and UDP. The exception here is the crystal structure of *Rickettsia bellii* (4HWG). We hypothesize that the cubic symmetry observed in this crystal structure may stabilize the closed conformation in the absence of a ligand, due to an increased number of interchain contacts. We also observe that (ii) in all cases where both substrate (UDP-GlcNac) and cofactor (UDP) are found bound in the crystal structure (1VGV, 3BEO, 4FKZ, 4NES, and 5DLD) the observed opening angle is reduced to 24-30°, significantly reduced as compared to the unbound state for MnaA. Finally, (iii) SaMnaA (5ENZ) and 1F6D are bound to UDP only. Here, both proteins have different opening angles for each monomer, leaving opening questions for this scenario. A possible explanation is that UDP could promote a variety of intermediate states going from a closed state with an opening angle of 26-27° up to more opened states with angles of 30-40°.

**Table 2.**
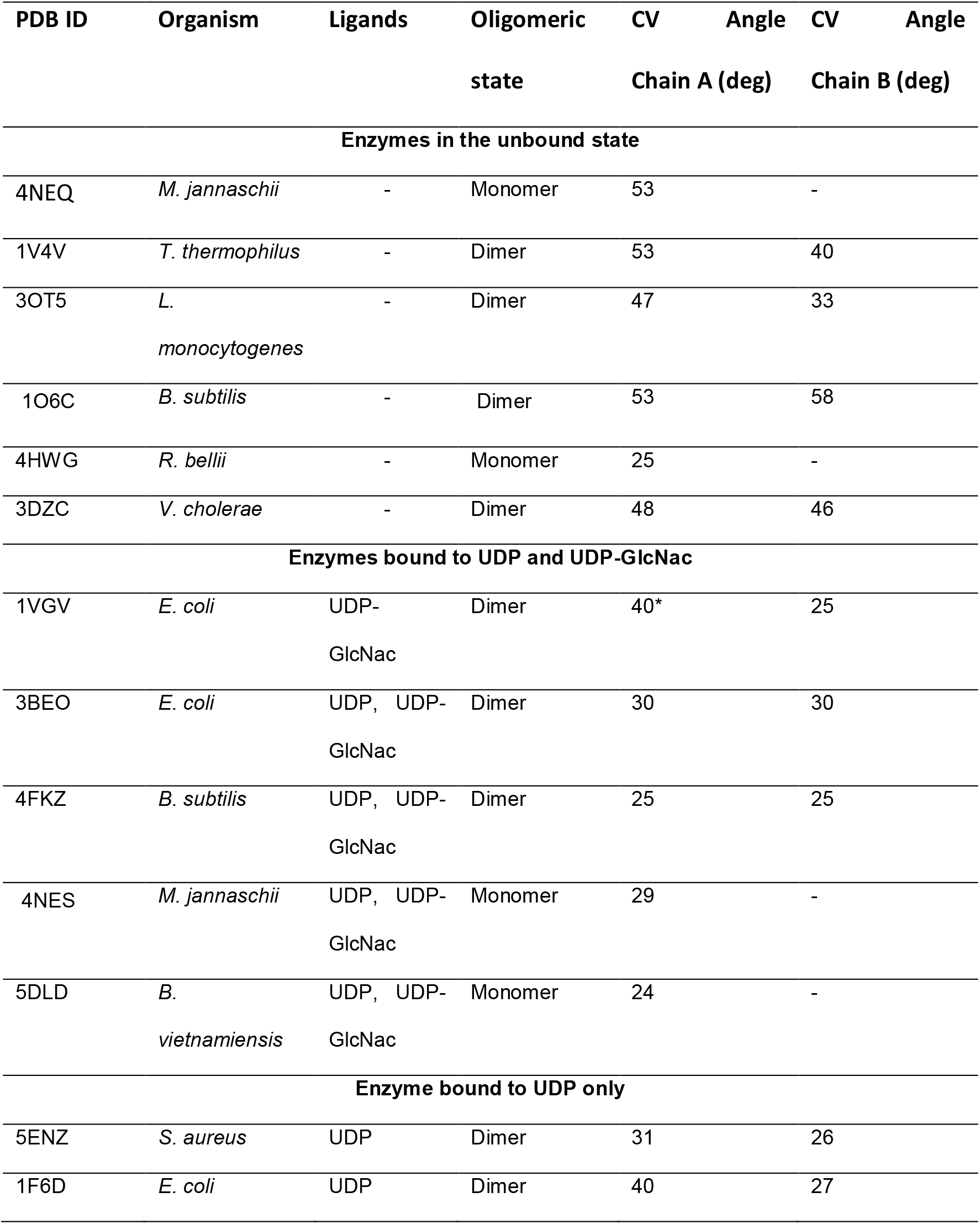
Opening state of known crystal structures of UDP-GlcNac 2-epimerases. *chain without ligands.

Altogether, the analysis of the available structural data indicates the existence of a major structural change in the MnaA that seems to be associated with the binding of UDP/UDP-GlcNac. However, the crystal structures are single conformations of an ensemble of thermally accessible structures that could be sampled in solution. In order to get a more detailed perspective of this structural change, we used MD simulations to generate an ensemble of conformations in unbound (APO) and ligand-bound (HOLO) scenarios.

### Equilibrium Simulations

The APO SaMnaA enzyme was simulated for 1 μs. During this simulation, a clear domain movement was observed in the MD trajectory and its magnitude could be measured by the change in the opening angle chosen as the collective variable (CV). As shown in Figures 1A and 1B, the enzyme accessed the opened conformation with an opening angle of about 53 degrees and a more intermediate state with CV angle of about 40 degrees. The time evolution of the opening was very fast, with opening happening still during the equilibration. On the other hand, once the structure was opened, it remained opened (or in the intermediary state) during the entire simulation, never visiting the closed state over the 1 μs simulation time.

**Figure 1.**
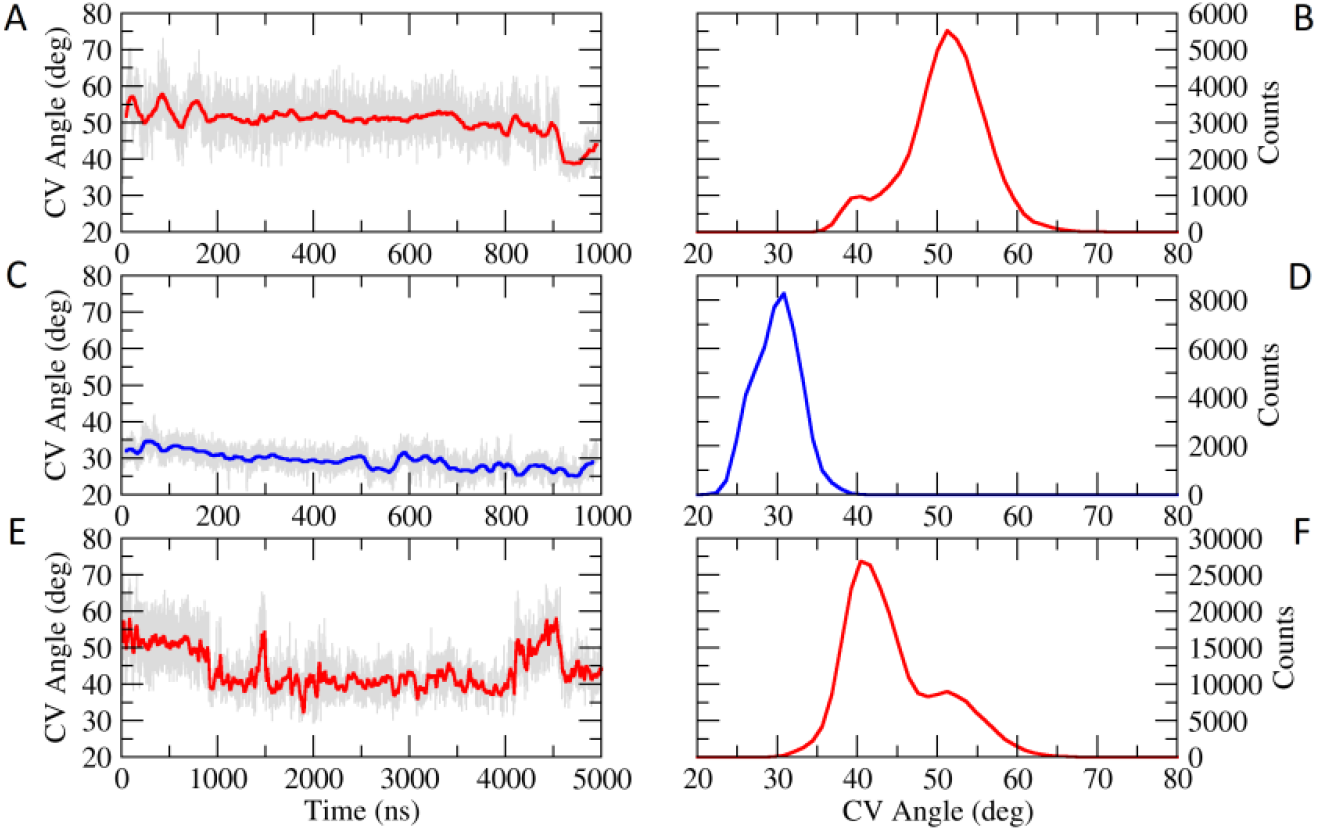
Equilibrium simulations of SaMnaA. (A) Opening angle observed during the simulation time for the 1-μs simulation of the APO SaMnaA structure. (B) Histogram of observed opening angle during the 1-μs simulation of the APO SaMnaA. (C) Opening angle observed during the simulation of the HOLO SaMnaA structure. (D) Histogram of observed opening angle in HOLO SaMnaA simulation. (E) Opening angle observed during the 5-μs simulation of the APO SaMnaA structure. (F) Histogram of the observed opening angle during the 5-μs APO SaMnaA simulation.

In order to increase the sampling, the initial 1 μs simulation was further extended up to 5 μs. Even in this longer timescale, the closed conformation was never visited (Figure 1E and 1F). Here, the intermediary state was mostly populated, resulting in a bimodal distribution of the CV angle with peaks at about 40 and 53 degrees, as shown in Figure 1F.

The 1-μs MD trajectory was analyzed using Principal Component Analysis (PCA) (Roe and Cheatham, 2013), and it was observed that the first two components described an opening movement of the enzyme domains (first PC) coupled to a torsion between them (second PC), as indicated by the arrows in Figure 2. Confirming the observations made based on the CV angle and the visual inspection of the trajectories, SaMnaA has a low-frequency domain movement towards an opened state in the absence of a ligand and cofactor.

**Figure 2.**
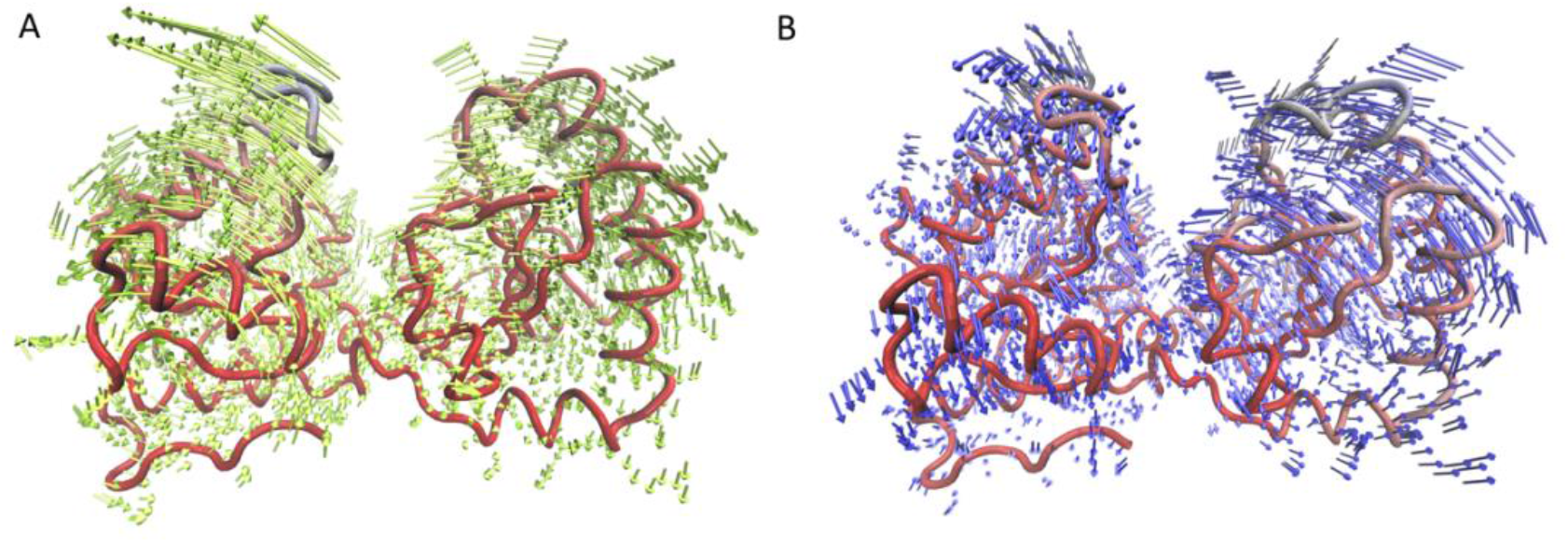
PCA Analysis of apo SaMnaA equilibrium simulation. Projections of the first (A) and second (B) components of the PCA analysis of the APO SaMnaA system MD trajectory.

The 5-μs MD trajectory of the APO system was further analyzed using PyEMMA (Scherer et al., 2015). Within this package, the MD trajectory (reduced to the representation of Cα atoms only) was clustered to define three major macrostates, as shown in Figure 3B. The representative structures show that the structures are going from an intermediate state to the opened one (Figure 3A and 3C). The first time-independent component (TIC), that describes the slowest movement of the system is in agreement with the behavior of the CV angle chosen, which indicates that this was a good choice of the CV indeed (Figure 3A). The Markov-State Modeling of the clusters allowed us to estimate a typical time for the closing movement, i.e., transitioning from the opened to the intermediary state, as 879 ns, and for the opening, i.e., going from the intermediary state to the opened state, as 423 ns. Since the closed state was never sampled in the productive simulation, it was not possible to estimate the typical timescale for the full opening of the enzyme.

**Figure 3.**
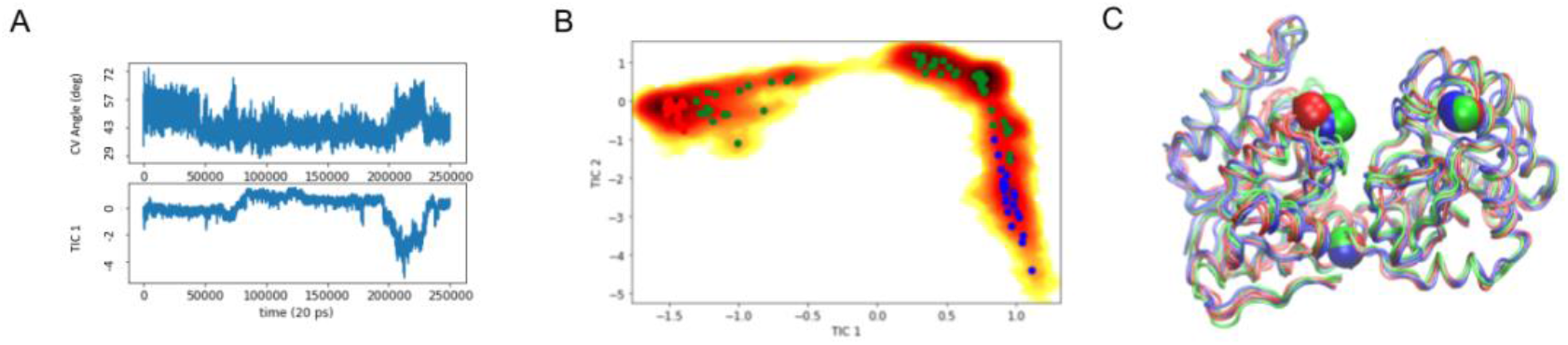
PyEmma Analysis of the 5 μs simulation of apo SaMnaA. (A) Comparison between the CV angle measured (upper plot) and the TIC 1 from the TICA analysis (lower plot). (B) Heatmap of TIC1 versus TIC2 where the systems stated were coarse-grained, resulting in the separation of three macrostates, shown here as cluster defined by colors red, green and blue. (C) Representative structures of the three states found for the system, with atoms used for CV definition atoms in evidence.

In summary, the results obtained from MD simulation for the APO state suggest that the unbound enzyme is found primarily in the opened state, but also sampling an intermediary state (very similar in structure to the opened state). The closed state was never observed, even in longer simulation times, consistently with the predominance of opened conformations observed in crystal structures available in the PDB.

The equilibrium molecular dynamics for the HOLO system showed a markedly different behavior as compared with the APO one (Figure 1C and 1D). Here, the enzyme shows a very well-defined preference for the closed state, with a typical CV angle of about 28-30 degrees. In agreement with crystal structures, the presence of the substrate and of the cofactor drives the equilibrium towards a closed conformation that is maintained during the entire simulation time. The trajectory was also analyzed using PCA (Figure 4) and the two first components represented on the protein structure do not show a clear coordinated movement towards the domain movement for opening, as observed for the unbound SaMnaA.

**Figure 4.**
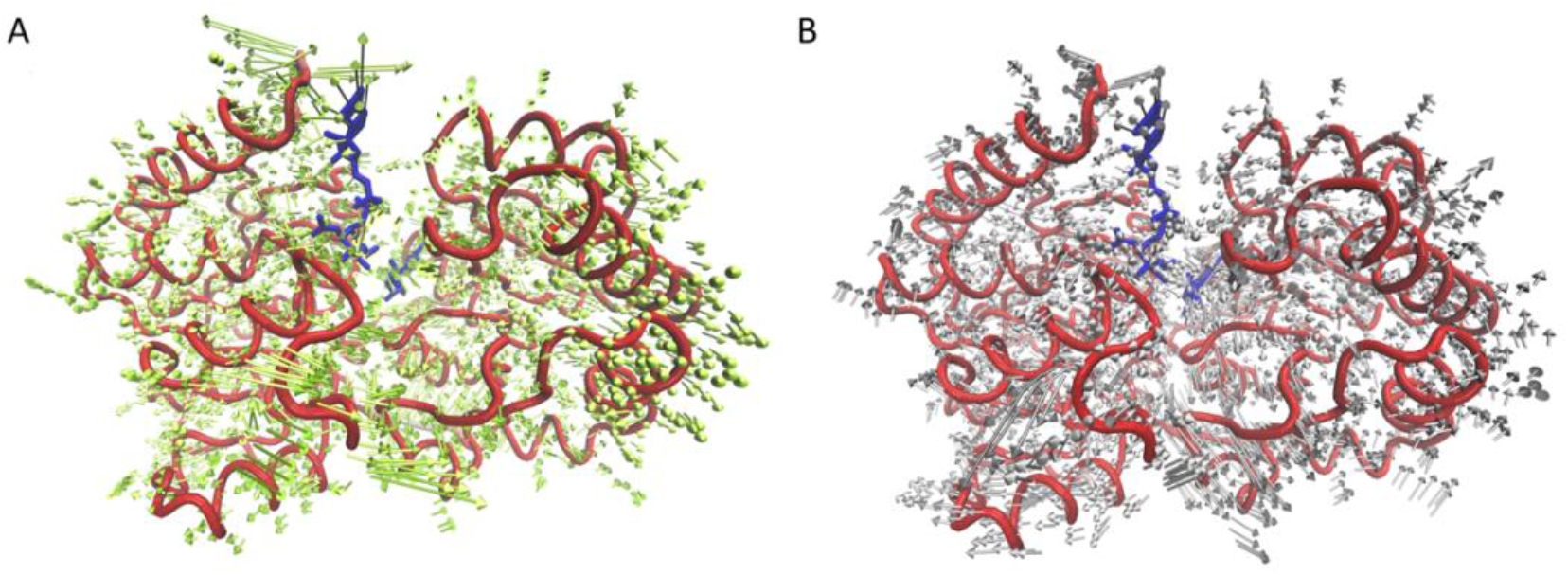
PCA Analysis of holo SaMnaA. Projections of the first (A) and second (B) components of the PCA analysis for the HOLO SaMnaA system MD trajectory.

For the HOLO SaMnaA, an analysis of the interaction energies involved in protein-UDP and protein-UDP-GlcNac interaction revealed a complex network of polar interactions involving the ligands, the enzyme and solvent atoms. Figure 5A shows a LIGPLOT (Laskowski and Swindells, 2011) representation of the interactions involving UDP for a snapshot of the equilibrium simulation. The interactions between UDP and the enzyme mainly involve the residues Glu288, Arg206, Ser282 and Arg11, in addition to several solvent molecules and an interaction between UDP and UDP-GlcNac. The individual contributions of these residues to the total interaction potential energy are shown in Figure 5C, where Arg11, and Arg206 stand out as major contributors due to their interactions with the phosphate groups of UDP. It is also interesting to note the positive (unfavorable) contribution from Glu288 for the UDP interaction (Figure 5C), due to the charge-charge repulsion with the UDP phosphate groups.

**Figure 5.**
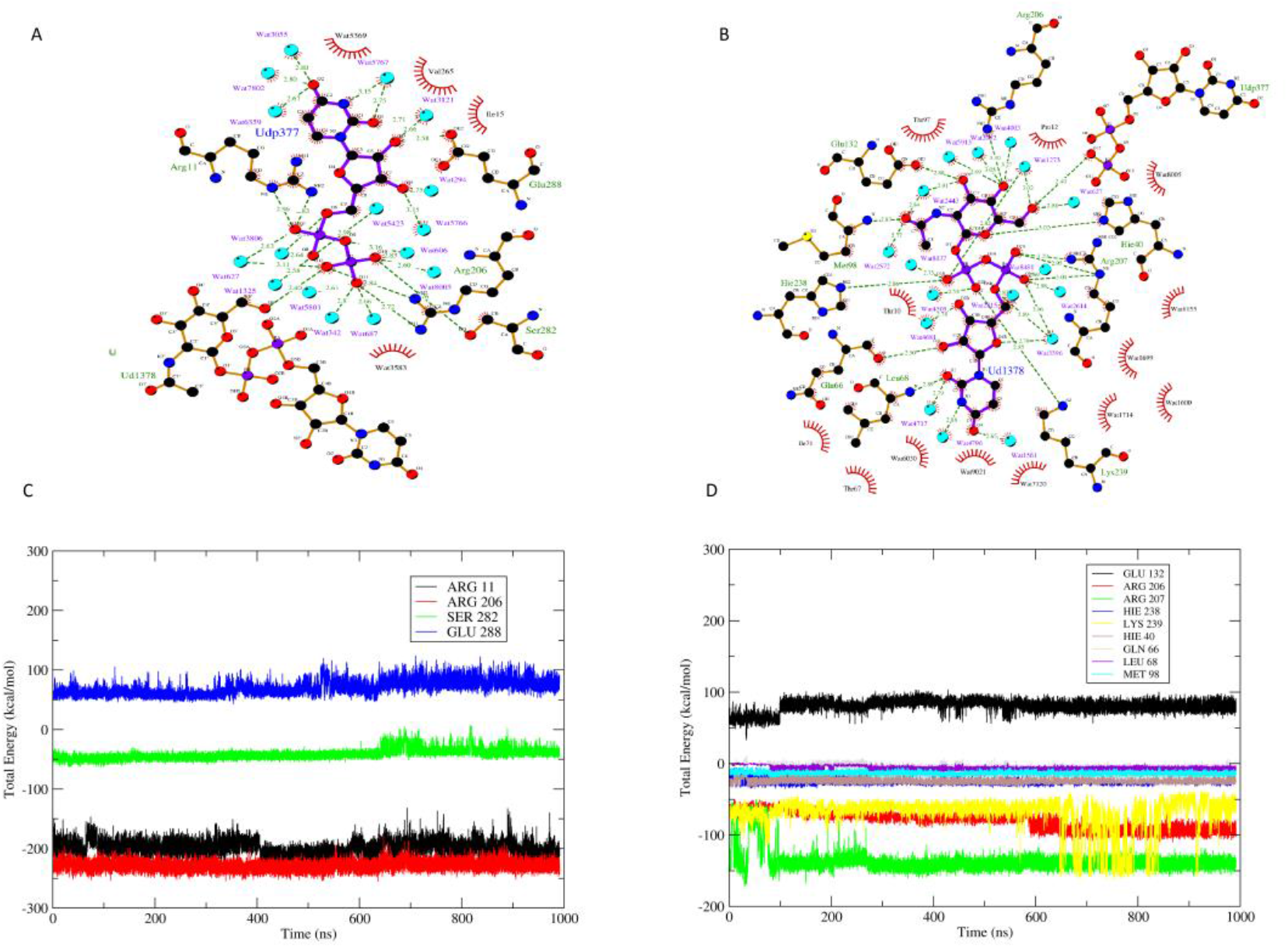
Analysis of UDP-N-acetylglucosamine and UDP interaction with SaMnaA. LIGPLOT depiction of the interactions between UDP (A) and UDP-GlcNac (B) with SaMnaA. Total interaction (potential) energy between UDP (C) and UDP-GlcNac (D) and its interacting amino acids are also shown.

For UDP-GlcNac (Figure 5B), the set of interacting residues include Arg206, His40, Arg207, Lys239, His238 and Glu132 in addition to the interactions with Leu68, Gln66, and Met98 through their main chain atoms. Again, as expected, the major contributors to the total interaction potential are the positively charged residues Arg206, Arg207 and Lys239, followed by the histidine residues His40 and His238 (Figure 5D). In conclusion, the interaction with UDP and UDP-GlcNac is maintained mainly through polar residues with the special contribution of positively charged residues in the vicinity of the active site.

### Energy Landscape of the Opening for the Unbound Enzyme

In order to better understand the energy landscape associated with the conformational changes coupled to substrate binding, we also run non-equilibrium simulations of the APO and HOLO systems. Although GAMD simulations generate noisier free energy profiles (potential of mean forces) when compared to umbrella sampling techniques, it does not require a pre-chosen CV. Considering this advantage, we used GAMD to probe the energy landscape of the domain movement in SaMnaA.

Three independent simulations of the APO system were performed with three different starting conformations. These conformations are three different coordinates extracted from the equilibrium simulation (0 ns, 50 ns and 100 ns of the productive simulation). Each conformation was simulated for 500 ns. In the accelerated simulations, we observed that in two simulations where the starting conformations with opening angles were close to 55° were reduced to intermediary states with opening angles close to 40-43°. In the third simulation, the enzyme evolved more flexibly, with the opening angle continuously varying between 45 to 65°, as shown in Figure 6A. The histogram of the accelerated simulations indicates a clear preference of the system for the intermediary state (Figure 6B). Interestingly, even with the boost potential, the APO enzyme never visits the closed state, suggesting a high energy barrier to reach this conformation for the unbound enzyme. We note that the PMF profiles observed in GAMD and ABMD simulations (Figure 6 and Figure 8) agree in general but are not similar. These differences are somewhat expected, since the methodologies rely on different assumptions. In GAMD, the simulation is accelerated by the introduction of boost potentials that do not consider the ‘reaction coordinate’ of the molecular event under evaluation, while in ABMD, the biasing potential is directed to this reaction coordinate. The final effect is a rougher, but bias-free PMF for GAMD and a finer, though biased PMF for ABMD. In this context, having both experiments set may be a good strategy.

**Figure 6.**
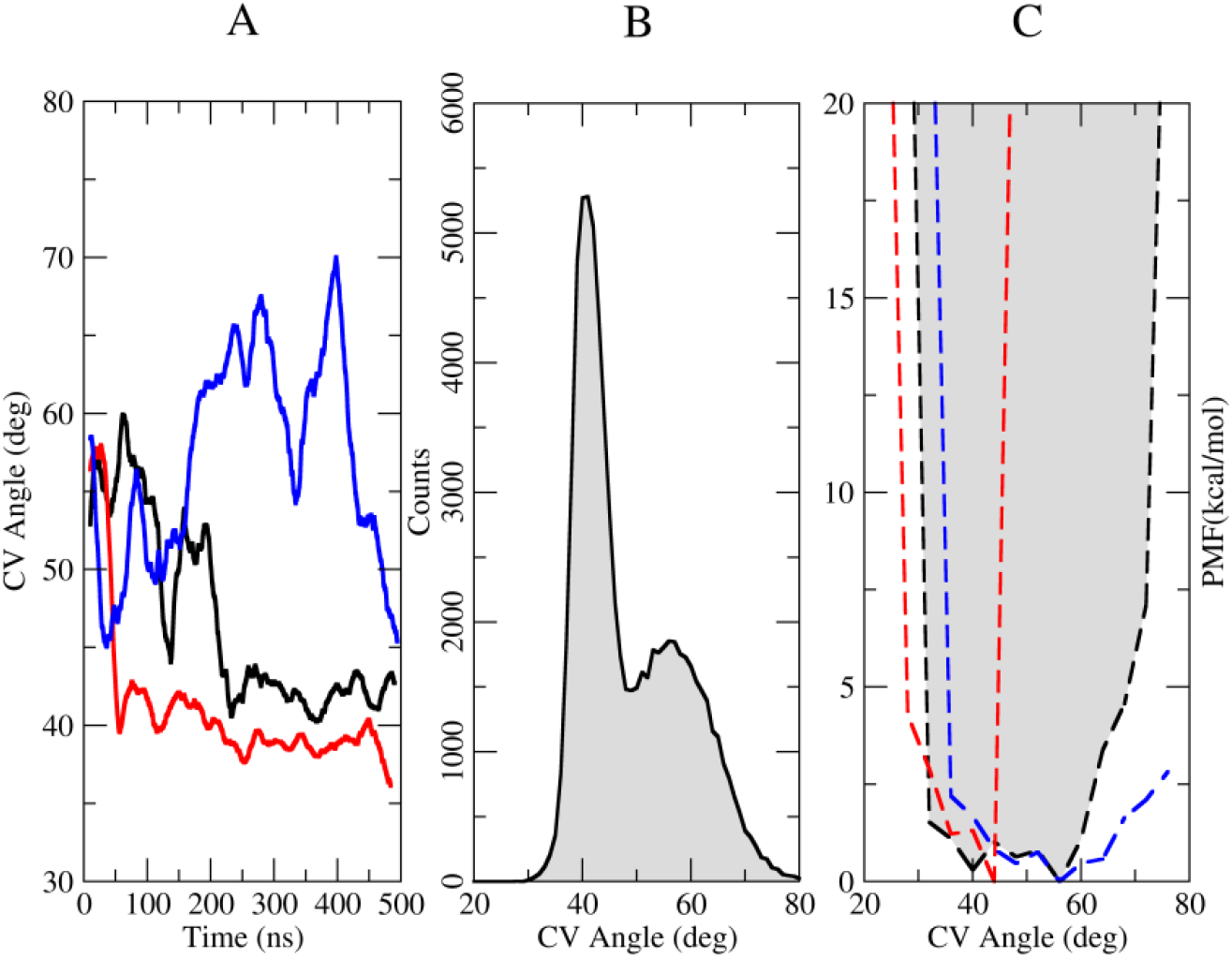
GAMD simulation for APO SaMnaA structure. The results from three simulations are shown: GAMD-APO-I in black curve, GAMD-APO-II in red curve and GAMD-APO-III in blue curve. (A) Observed CV angle versus simulation time. (B) Distribution of observed CV angles during the simulation. C) PMF computed after reweighting.

Potential of mean force (PMF) profiles were computed from the GAMD simulations and are shown in Figure 6C. Two slightly different profiles can be traced from the computed PMFs: (i) in all cases, the energies increase rapidly when the opening angle is below 35°. The simulation GAMD-APO-I (black curves in Figures 6A and 6C) shows a low energy for the conformations encompassing opening angles between 35 to 55 degrees, GAMD-APO-II (red curves in Figures 6A and 6C) shows low energies associated with opening angles between 35 to 45 degrees and GAMD-APO-III (blue curves) show low energies associated with opening angles between 35 to 70 degrees. All simulations agree that the intermediary state is a low energy state for the apo system, and simulations I and III also suggest that the opened state (CV angle > 50°) can be thermally accessed by the unbound enzyme. Although the PMFs computed using GAMD data are coarse, the overall results agree with the 5-μs simulation data, suggesting that, once unbound, a putative equilibrium between conformations with opening angles between 40-45 degrees (intermediate) and 50-60 degrees (opened conformation) are expected to be found in solution.

Similarly, three independent GAMD simulations for the HOLO SaMnaA were conducted and analyzed. The simulations also showed that the bound protein tends to stay in a closed conformation, with an opening angle of about 30 degrees, as shown in Figure 7A. Here, two simulations remained the entire simulation time in the closed conformation (GAMD-HOLO-II and GAMD-HOLO-III, shown in Figure 7A in red and blue curves, respectively) while the other simulation (GAMD-HOLO-I, shown in black curve in Figure 7A) showed a continuously opening angle, starting with about 26 degrees at the beginning of the GAMD simulation and reaching about 48 degrees at the end of the simulation. Interestingly, in this simulation, the ligand (UDP-GlcNac) was observed to unbind from the enzyme at a simulation time of 175 ns, as indicated in the binding energies and RMSD computed from this trajectory (Figures 7D and 7E).

**Figure 7.**
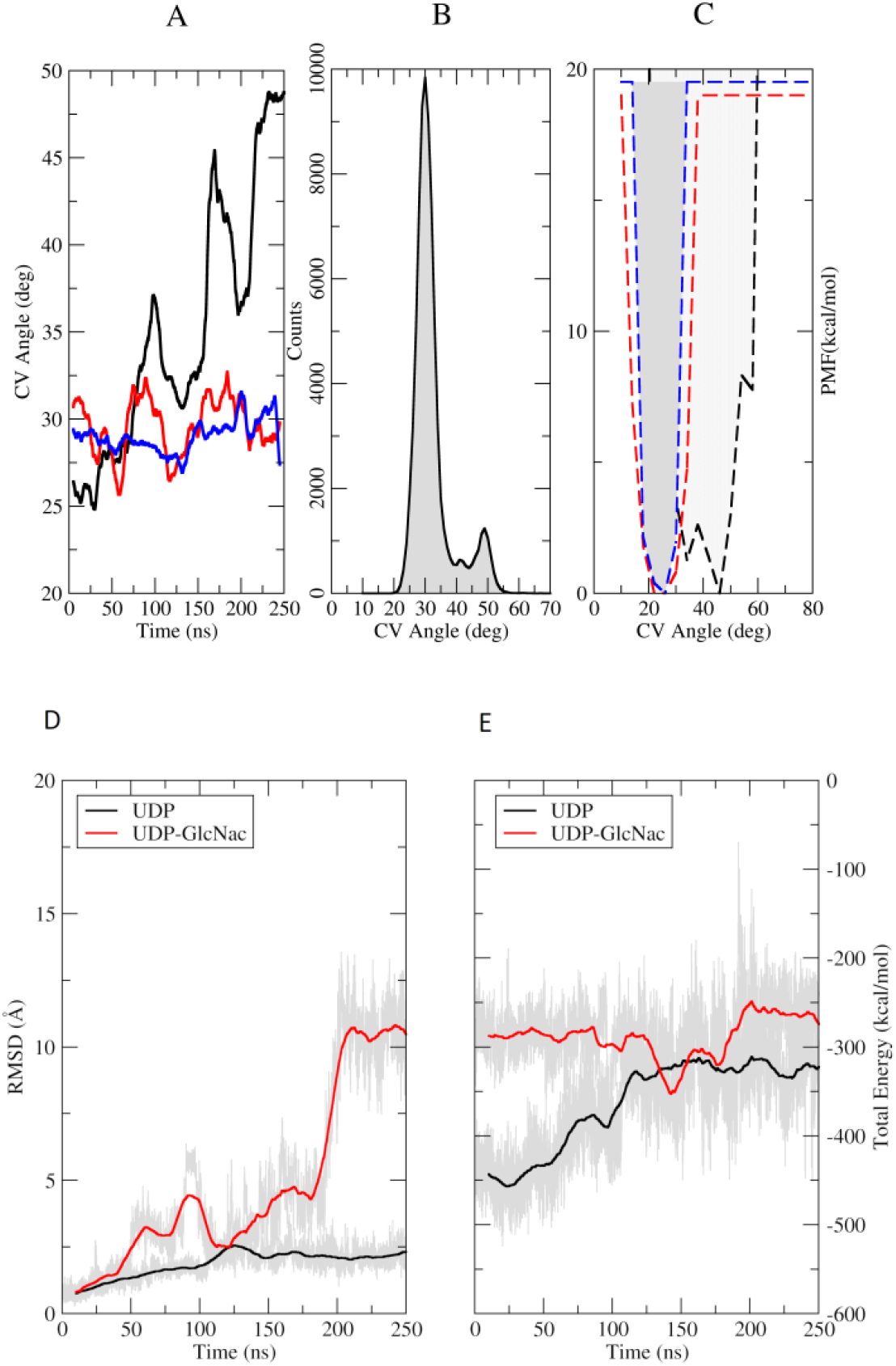
GAMD simulation data for the HOLO SaMnaA. The results from the three independent simulations are shown: GAMD-HOLO-I in black curve, GAMD-HOLO-II in red and GAMD-HOLO-III in blue curve. (A) Observed CV angles during the simulation time. (B) Histogram of the observed angles during the simulation. (C) Computed PMFs from the energy reweighting. (D) RMSD calculated for the UDP (black) and UDP-GlcNac (red). (E) Total interaction energies computed for UDP (black) and UDP-GlcNac (red) with the protein.

The PMFs calculated for the GAMD simulations of the HOLO system (Figure 7C) indicated a clear minimum in the free energy associated with an opening angle of about 25 degrees. For the simulations GAMD-HOLO-I, where the unbinding event was observed, a shift in the minimum towards the intermediary state was observed, consistent with minimum expected for an APO enzyme. The unbinding of the UDP-GlcNac causes a major shift in the UDP position inside the active site, forcing it to lose some of the interactions with the protein. Altogether, the histogram of the opening angle over the three GAMD simulations clearly indicates a preference for the closed state for the bound system, consistently with the equilibrium simulations. In addition, the unbinding event observed due to the acceleration applied in our non-equilibrium simulation clearly suggests that the opening event is coupled with the absence of the ligand.

The coarse description of the energy profile associated with the conformational state of SaMnaA may be associated with the poor sampling of the transition between the states, which is rarely observed in our simulations. In this way, a finer energetic description can be obtained for enhanced sampling techniques where the reaction coordinate or collective variable is systematically sampled over the simulation. Looking for this finer description of the opening/closing movement in SaMnaA, we explored the enzyme dynamics with Adaptatively-Biased MD (ABMD).

Three independent ABMD simulations were conducted for the APO SaMnaA, with the same starting coordinates as chosen for GAMD. Each simulation explored the opening angle as a CV with a range of 10-100 degrees and resolution of 1 degree in 250 ns of ABMD simulations. The results obtained, shown in Figure 8, indicate that the CV angles in the range of 40-60° lie in the minimum of the free energy landscape, with energy barriers of about 10 kcal/mol (~16 kT) to reach a closed state, defined as a CV angle of about 25 degrees. The first simulation (ABMD-APO-I) showed an interesting broad minimum covering the entire region defined as the intermediary and opened state, while simulations ABMD-APO-II and ABMD-APO-III showed more defined minimum at about 55°, consistent with an opened state. Together, these simulations indicate a clear energy minimum covering opened or intermediary-to-opened state for the unbound enzyme.

**Figure 8.**
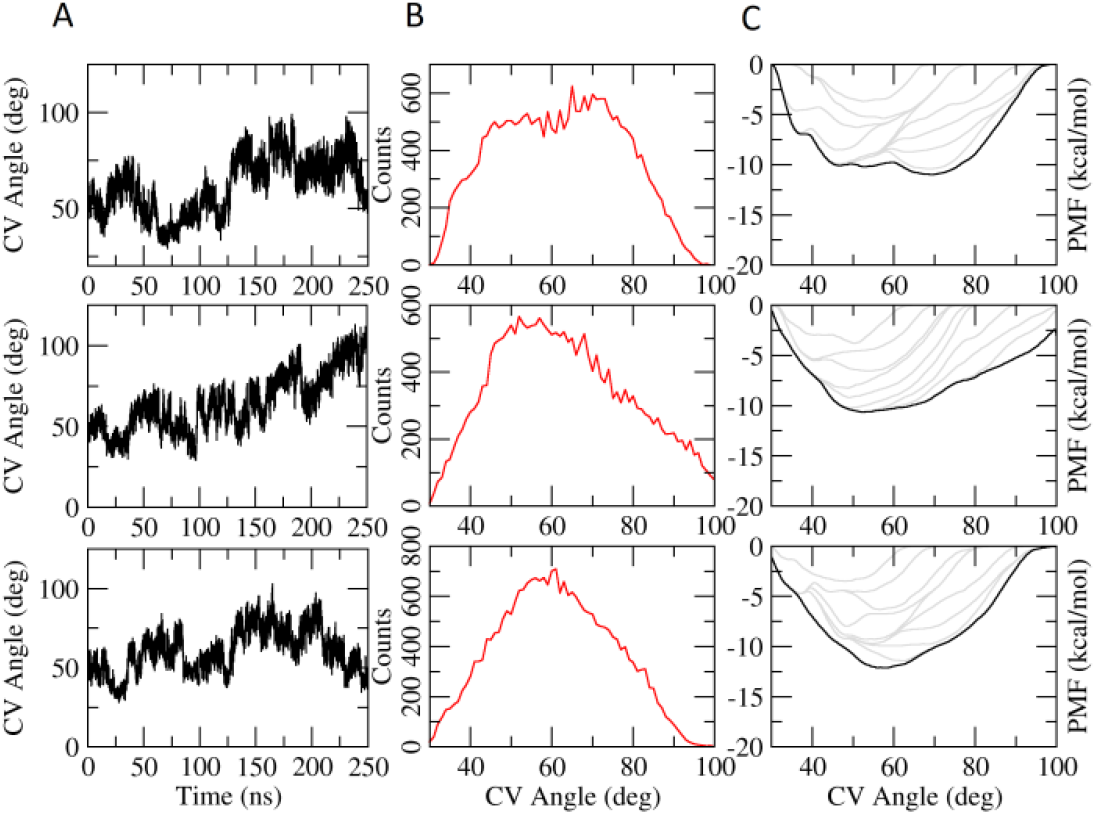
ABMD of the APO system for simulations ABMD-APO-I (upper row), ABMD-APO-II (middle row) and ABMD-APO-III (bottom row). (A) Time evolution of the CV angle; (B) CV angle distribution; C) Potential of mean force. The gray lines indicate the time evolution (and convergence) of the PMF in 10 step ABMD simulations of 25 ns each.

For the HOLO system, ABMD simulations accessed a wide range of CV angles, including the opened state. Again, once forced towards higher CV angles, the substrate was dissociated from the enzyme structure, resulting in the unbound substrate and the opened enzyme. The correlation between unbinding and opening becomes clear when Figure 9 is compared with Figure S1 (supplementary material), that shows the interaction energy between the enzyme and substrate. The simulation data indicate that the presence of the ligand is crucial for holding the closed conformation stable and, once the conformation is destabilized (by external forces, in this case), the ligand is spontaneoulsy dissociated from the enzyme.

**Figure 9.**
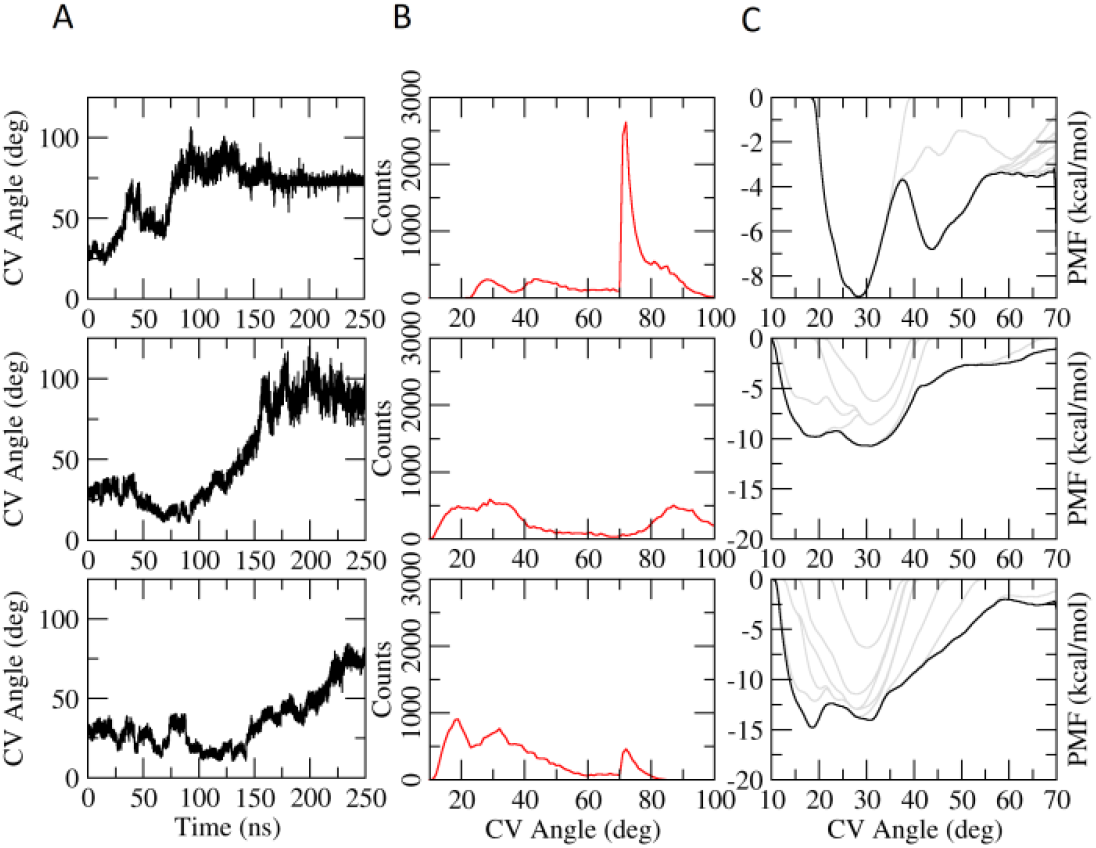
ABMD of the HOLO system for simulations ABMD-HOLO-I (upper row), ABMD-HOLO-II (middle row) and ABMD-HOLO-III (bottom row). (A) CV angle evolution during the simulation. (B) Histogram of CV angle. (C) Potential of mean force computed for the ABMD simulation data. The gray lines indicate the time evolution (and convergence) of the PMF in 10 step ABMD simulations of 25 ns each.

An important question remains unanswered: what is the mechanism involved in ligand binding and enzyme domain movement? Or, what is the nature of the driving force for the domain movement observed in the presence of a substrate? A visual analysis of the enzyme structure reveals an extensive network of positively charged amino acids in the vicinity of the substrate pocket. These residues stabilize the interaction with the phosphate groups of the UDP-GlcNac, thus making the electrostatic contribution of the binding energy as the largest contribution for binding. In the other hand, in the absence of a substrate, or product, the interaction of these positively charged residues is very unfavorable and could be alleviated if the residues are moved apart in a domain movement. Thus, we hypothesized a major driving force for this large-scale movement may be the electrostatic energy associated with the interaction of positively charged residues.

In order to evaluate this hypothesis, the APO and HOLO structures were analyzed using the Delphi Force web server (Li et al., 2017) to assess the magnitude of electrostatic forces between the N-terminal and C-terminal domains of SaMnaA. For the HOLO enzyme, the ligands were set as part of the C-terminal domain. The computed net forces are represented as arrows in protein structure in Figure 10. Here we observe that, in the absence of the substrate/cofactor, a net force vector is observed as pointing towards the direction of the domain movement (orange arrow in Figure 10). In the presence of the substrate, the net force is shifted to the twisting movement between the domains (also observed in the equilibrium simulation for the HOLO system). So, the analysis of the electrostatic forces provides a piece of evidence favoring the electrostatic term as a major driving force for the domain movement.

**Figure 10.**
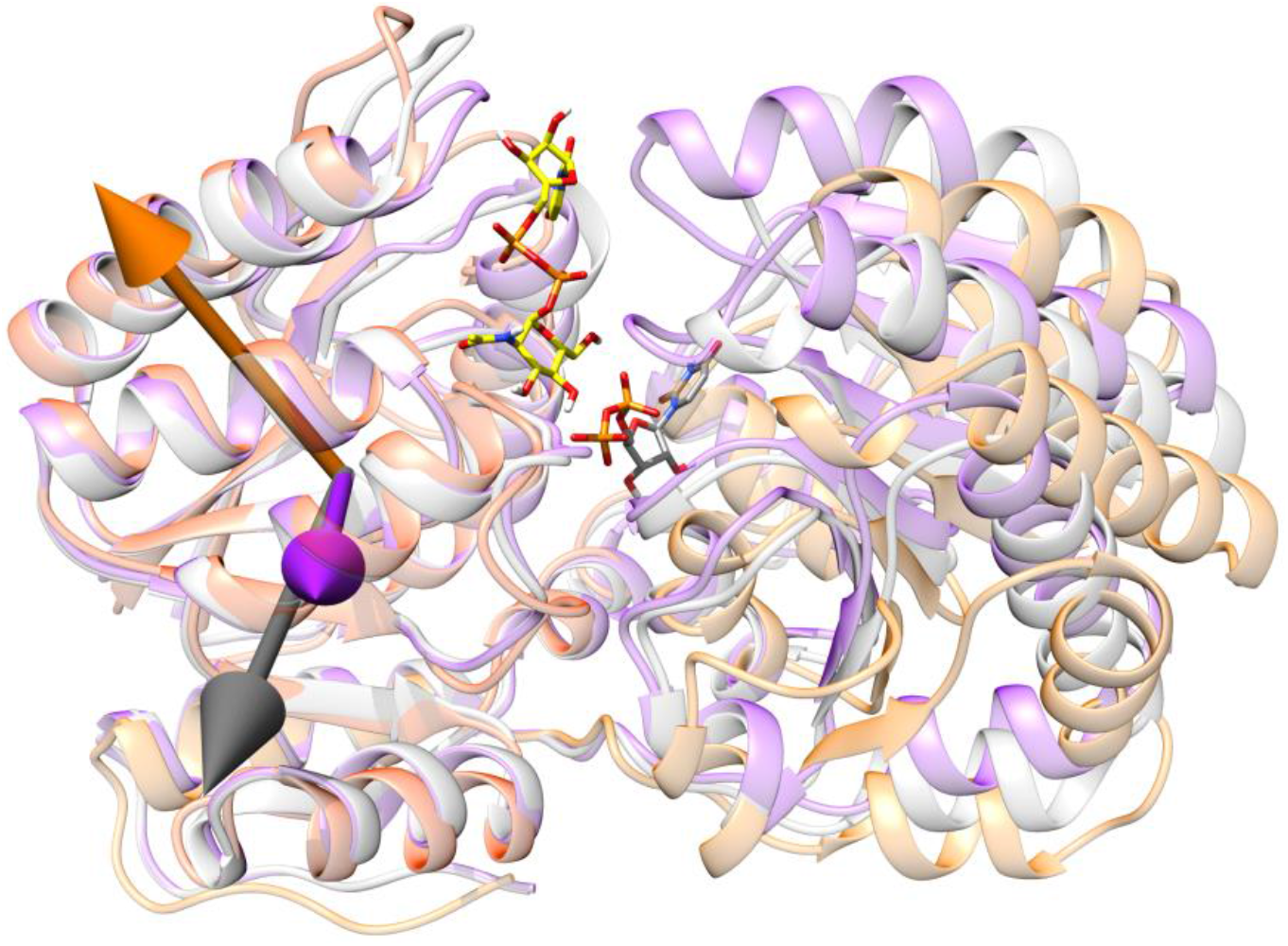
Electrostatic forces on SaMnaA structures. The sum of the resulting interaction electrostatic forces between the C-terminal and N-terminal domains for the open (orange), intermediary (gray) and closed (purple) states of the protein, with ligands in the active site (cyan).

## Discussion

In this work, we used over 10 μs of simulation data to evaluate the mechanisms involved in the structural rearrangement of *S. aureus* UDP-N-acetylglucosamine 2-epimerase as a function of the binding of the substrate. (Table 1). Our equilibrium and non-equilibrium simulation data showed that the enzyme is remarkably stabilized in a closed conformation when bound to substrate and cofactor, while the unbinding event leads to a change in the enzyme equilibrium structure towards a more opened structure. The analysis of the internal forces for the opened and closed conformation strongly suggested that the electrostatic term may be a major driving force for the enzyme opening, alleviating the unfavorable interactions between positively charged residues in the absence of the electron-rich substrate.

Considering the electrostatic forces as the main molecular driving forces acting to drive the ligand-dependent conformational changes, several additional experiments could be designed to further explore these molecular driving forces. We extended our analysis of this molecular mechanism using two additional experimental evidenced. First, electrostatic forces should be captured by simpler molecular models, as such used in coarse-grained models. We used SIRAH force field (Darré et al., 2015; Machado et al., 2019) to generate a simpler (coarse-grained) model but still retaining most of the polar description of the protein. After 23 μs of equilibrium simulation, we observed the APO enzyme to only sample the opened conformation (Supplementary Figure S2) with a similar distribution of opening angle as observed in the atomistic simulation, suggesting that the molecular driving force that forces the apo enzyme to the opened conformation is kept even in this simpler model. As a second experimental evidence, we introduced set of mutations in enzyme surface to reduce the number of positively charged residues close to each other in the enzyme domains and/or to introduce ‘charged clamps’ by introducing negatively charged residues able to interact with positive residues in the opposite domain. As shown in Supplementary Figure S3, from the mutants evaluated, at least two mutants were shown to be very effective in maintaining the closed conformation even in the absence of a ligand, thus reinforcing the electrostatic force as a major driving force for the domain movement in SaMnaA.

The shift in the energy landscape, caused by the change in the environment (substrate in the active site) agrees with the previously proposed concept of a dynamic energy landscape (Kumar et al., 2000), which is an important feature for the protein function since the macromolecule can adjust its population accordingly to changes in the environment, such as pH shifts or the presence of substrates. Kumar and coworkers (Kumar et al., 2000) proposed two main types of dynamic energy landscapes. The Type I is characterized by an environment change that causes a shift in the energy minimum of the energy landscape, thus shifting the preferential conformation of the protein. Type II, on the other hand, describes the situation where the change in the environment causes a change of the roughness in the surface, where the conformations that were trapped in an energy minimum now have a lower barrier to overcome and change into another conformation. Within these definitions, the dynamic landscape observed for SaMnaA seems to be closer to type I, that is usually observed in enzymes that are allosterically regulated, or in multimolecular complexes.

As previously observed for other enzymes, the domain movement may also be an outstanding structural requirement for catalysis. The closed state, observed in the presence of the substrate, allows the active pocket to exclude solvent molecules, resulting in a low dielectric medium with higher magnitudes of electrostatic forces acting and finally allowing the catalytic reaction to proceed. Additionally, once the reaction is completed, the opening movement allows the diffusion of products and new substrate molecules (Kumar et al., 2000; Tsai et al., 1999). As far as these authors are aware, there is no experimental evidence of the mechanism of product scaping from the active site. By judging by the catalytic cleft structure, it seems plausible that the product could diffuse after a spontaneous opening of the enzyme, with UDP-ManNac scaping first and UDP remaining bound or scaping after UDP-ManNac. However, further investigations are still necessary to better elucidate this structural mechanism.

SaMnaA may be an important target for the development of drug candidates to fight resistant infections. The molecular simulations and previous data show that the ligands may enter the active site when the protein is in the open/intermediate conformation, being closed by the interactions made between the molecules and the protein. Also, positively charged residues are involved in the most energetically favorable interactions. This indicates that the open conformation should be targeted, and the electrostatic forces should be taken into account when screening for new drug candidates.

In conclusion, here we explored the dynamics of the *S. aureus* MnaA that plays an important role in the biosynthesis of the wall teichoic acid and in the cellular resistance to β-lactam antibiotics. The protein has an open/close movement, important for the maintenance of the substrate in the active site during the catalysis. This movement is mediated by the ligands, which interact with positive amino acids of the protein, shifting the energy profile for the system. The dynamic energy landscape for the systems APO and HOLO suggest that there is a population of preferential states that shifts with the change in the environment. Finally, a structure-based mechanism is proposed, providing a rationale for the domain movement observed for SaMnaA.

## Acknowledgements

We thank the fruitful contributions made by prof. Milton T. Sonoda (*in memorium*). The authors thank the financial support provided by Fundação de Amparo à Pesquisa do Estado de São Paulo (FAPESP) through grants 2017/18173-0, 2015/26722-8 and 2015/13684-0 and by Conselho Nacional de Desenvolvimento Científico e Tecnológico (CNPq), through grant # 303165/2018-9. This study was financed in part by the Coordenação de Aperfeiçoamento de Pessoal de Nível Superior - Brasil (CAPES) - Finance Code 001.

