## Supplementary Material for "Energy Landscape of the Domain Movement in *Staphylococcus aureus* UDP-N-acetylglucosamine 2-epimerase"

### Supplementary Figure S1

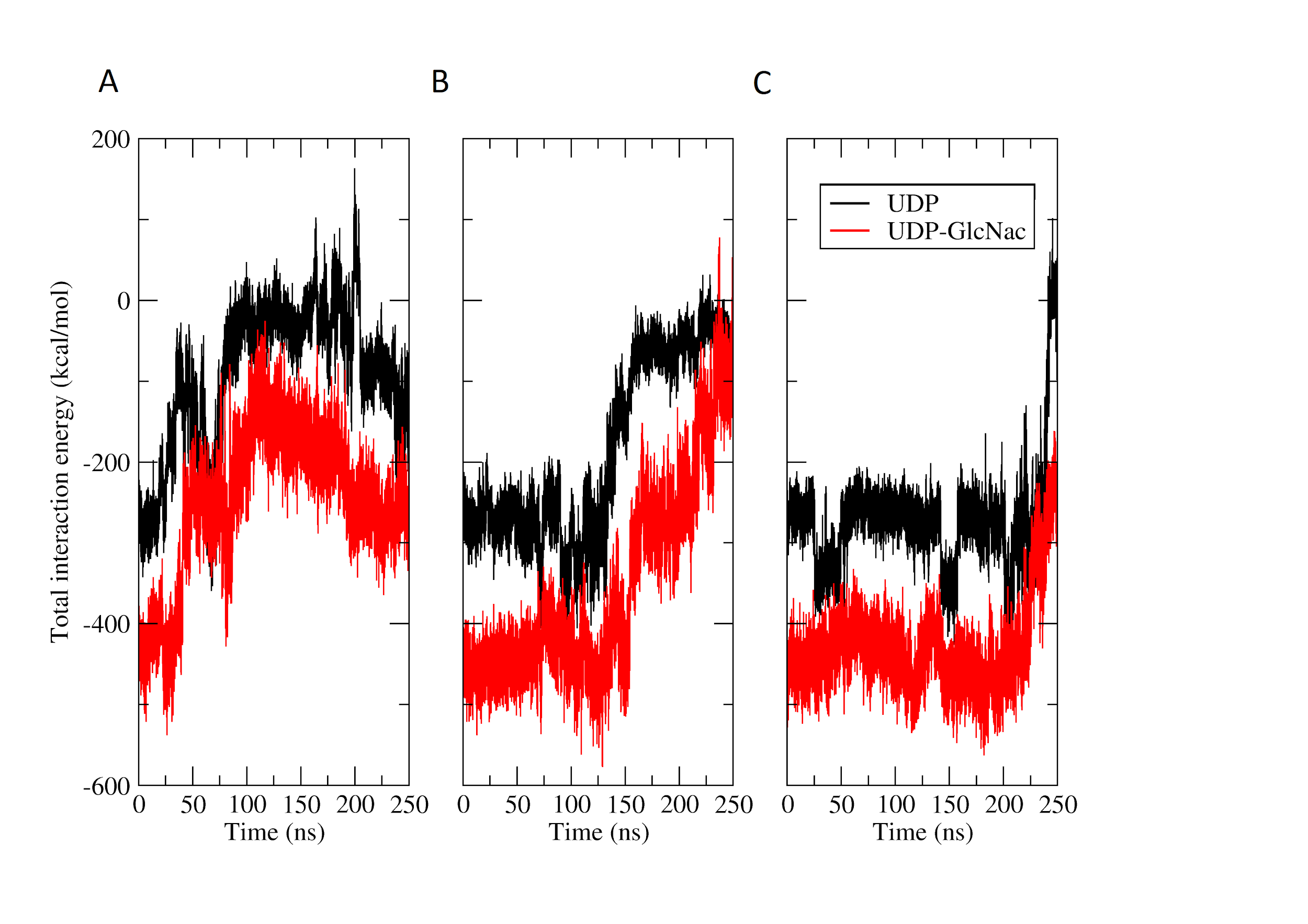

**Figure S1** - Total interaction (potential) energy computed between the UDP (black) and UDP-GlcNac (red) with SaMnaA for the three ABMD simulations.

### Supplementary Figure S2

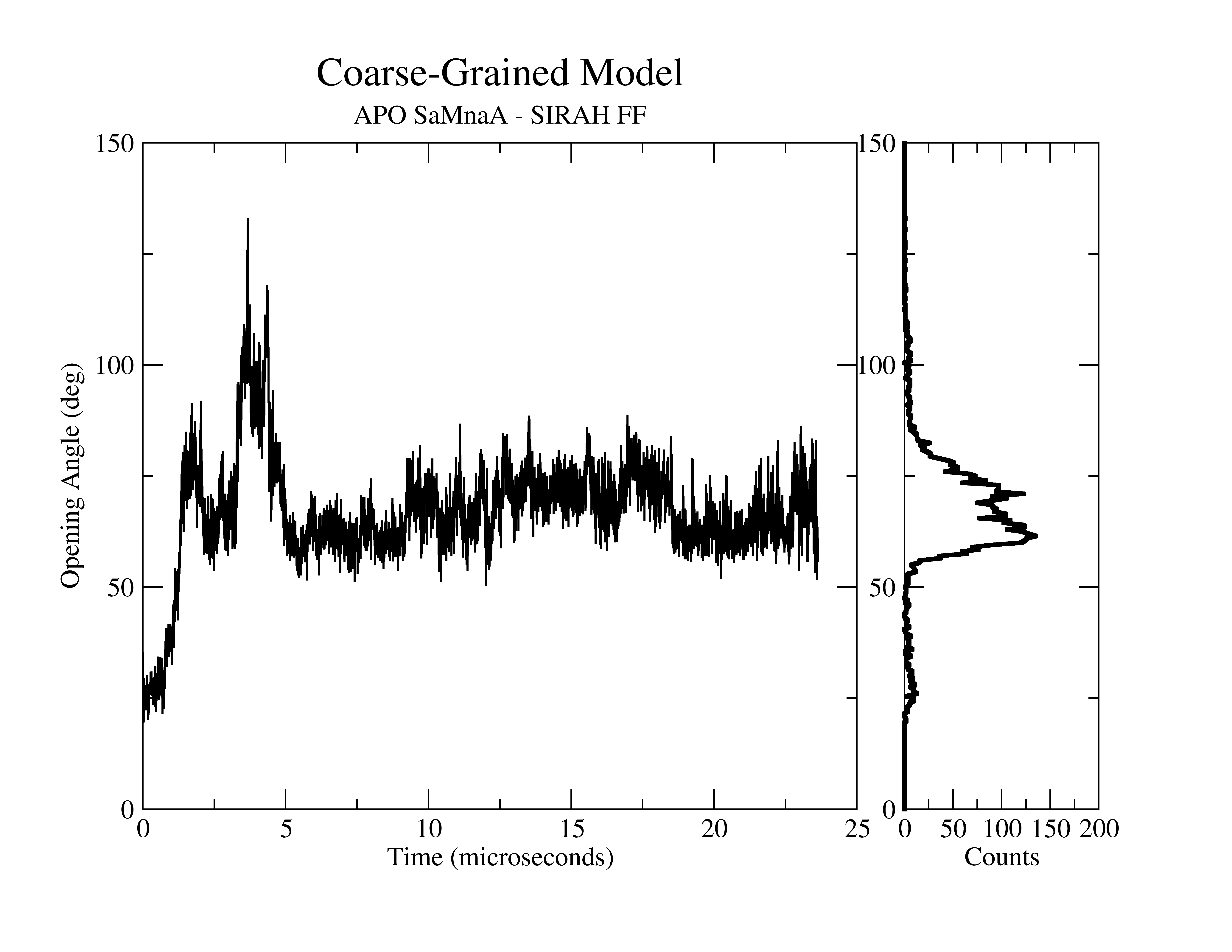

**Figure S2**. Time evolution of the opening angle in an equilibrium simulation of APO SaMnaA using a coarse-grained model parametrized with SIRAH force field (left). After 23 microseconds, a very similar sampling of conformational states was observed when compared to the fully atomistic simulation (Figure 1 of the main paper), suggesting that the major driving force for opening is kept by the simpler coarse-grained model. The right panel shows the histogram of the sampled states mapped on the opening angle.

### Supplementary Figure S3

| 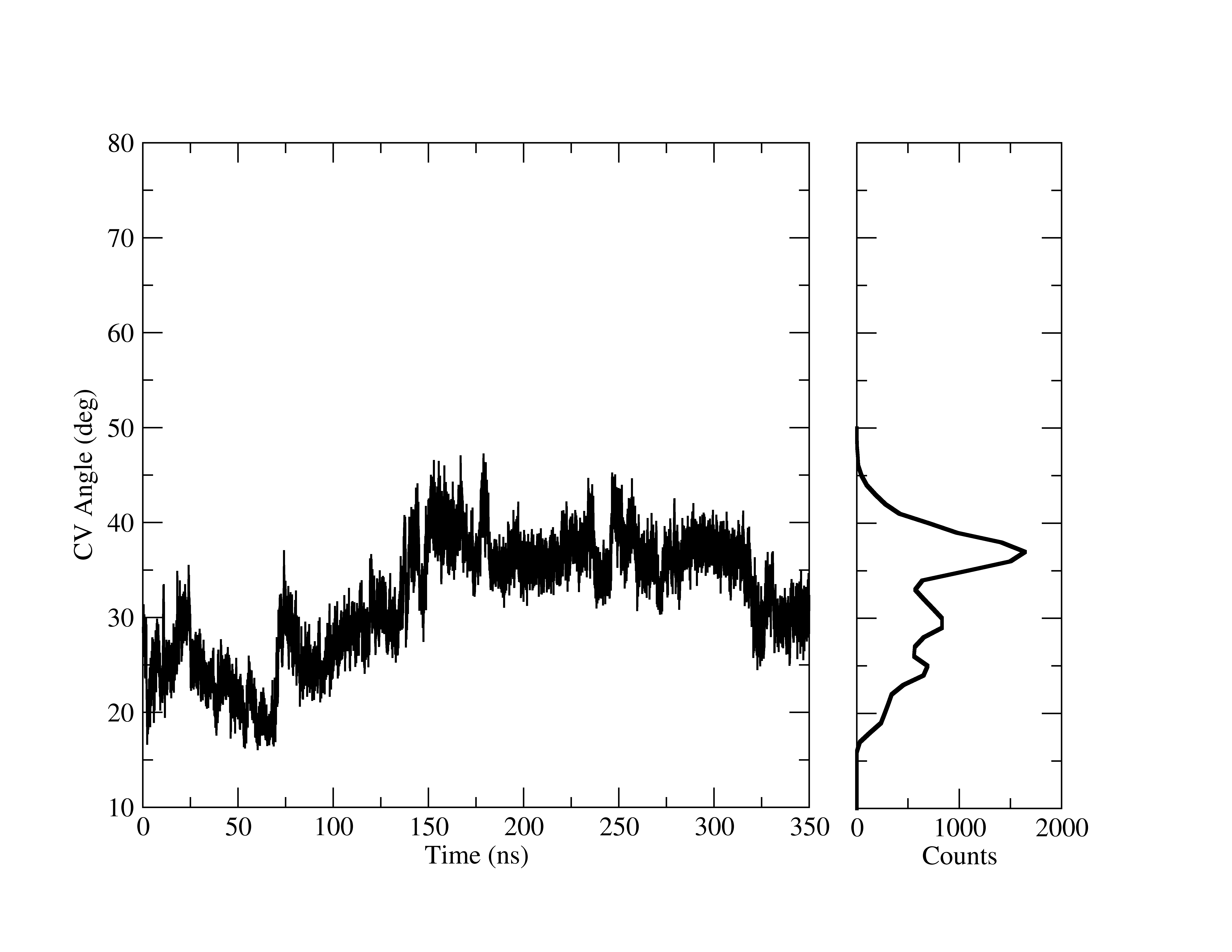 | 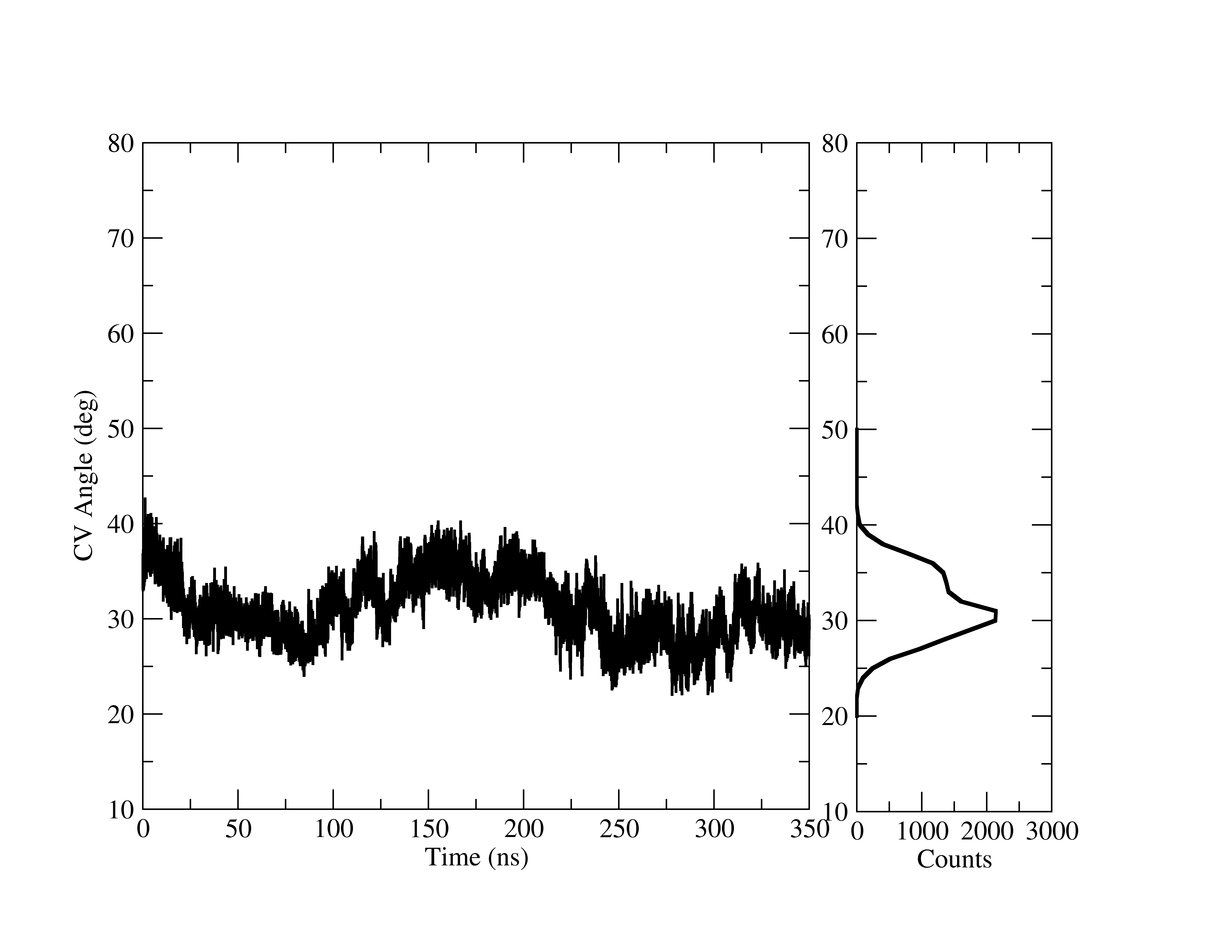 |
| --- | --- |
| 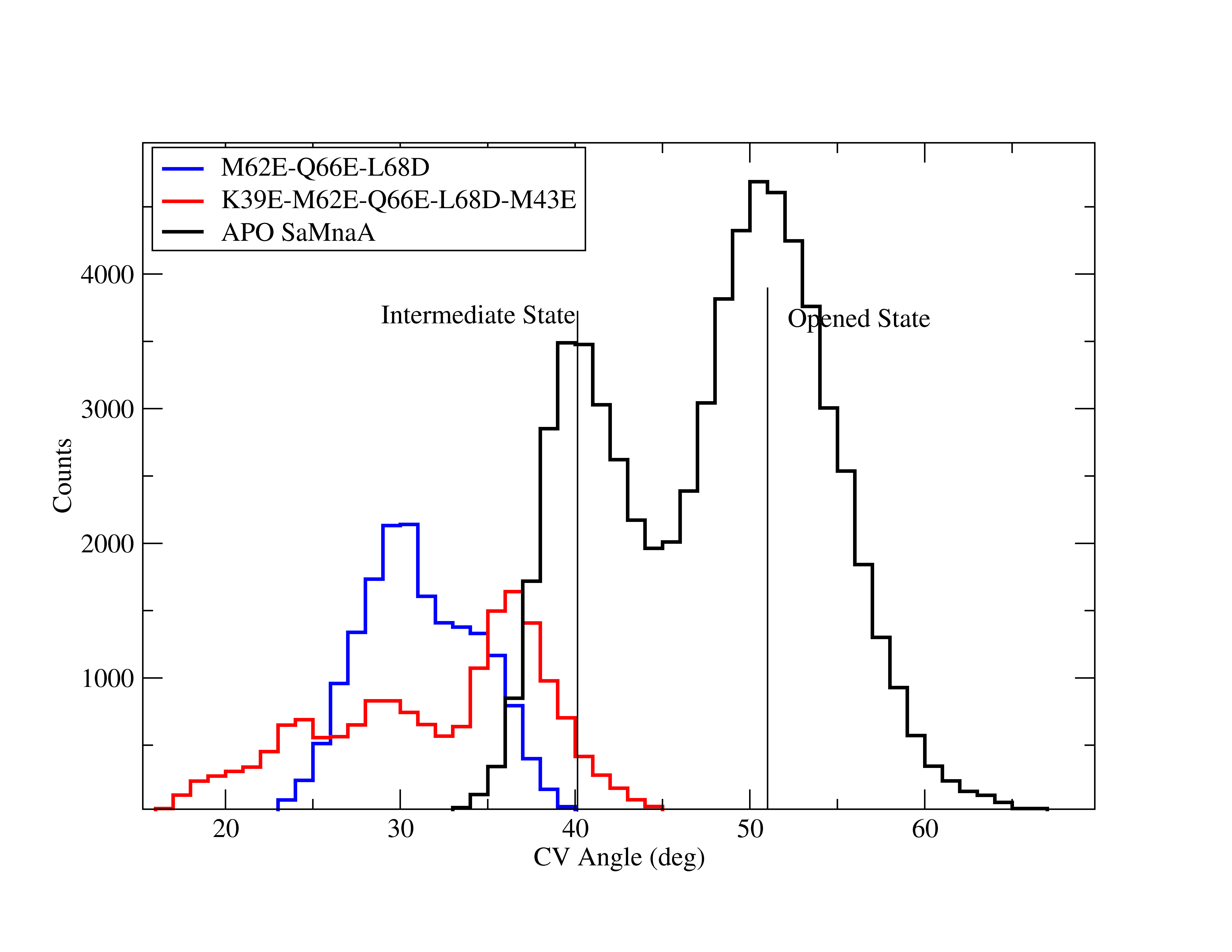 | 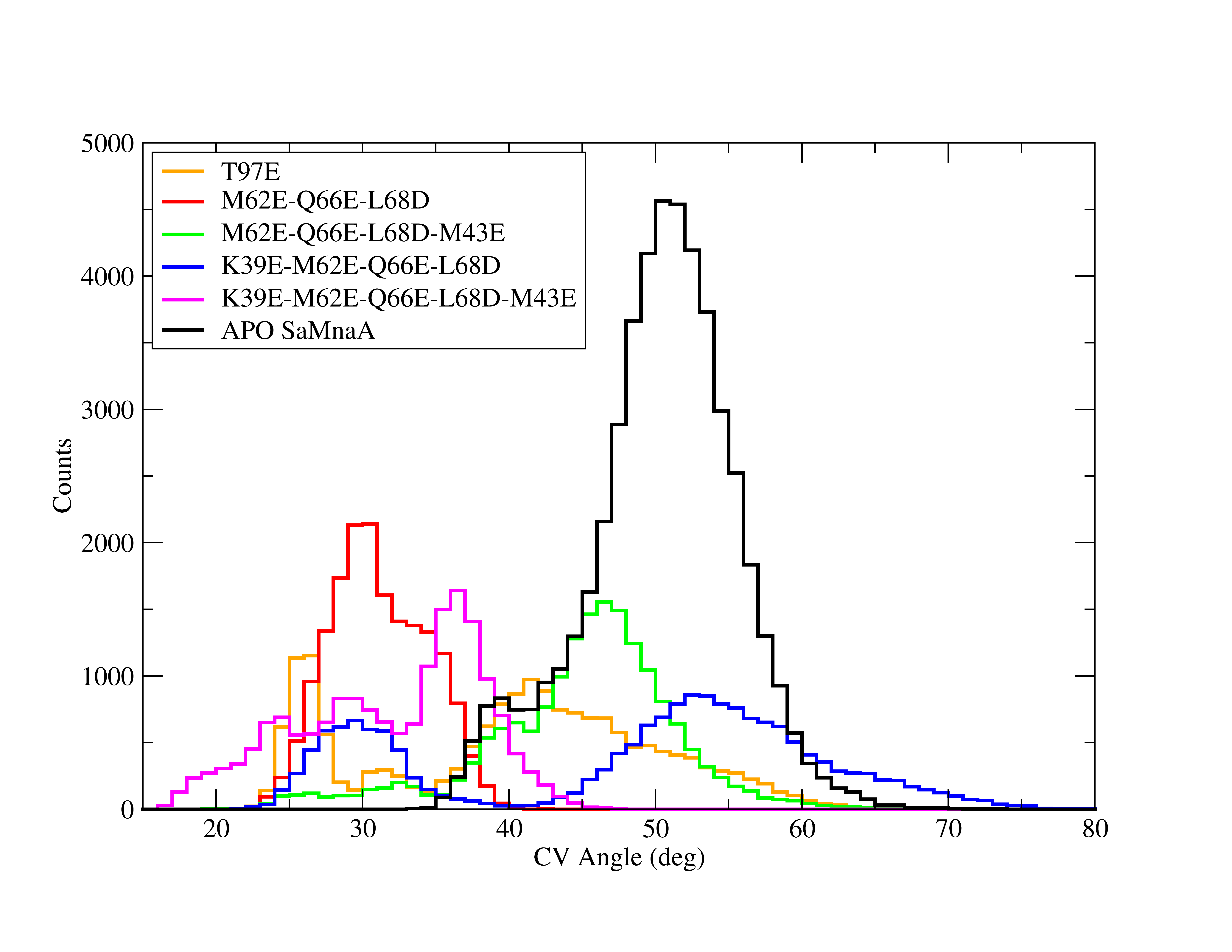 |

**Figure S3**. (*Top row*) Time evolution of the opening angle for two mutants: K39E-M62E-Q66E-L68D-M43E (left) and M62E-Q66E-L68D (right). (*Bottom left*) Histogram of opening angles sampled in the APO SaMnaA (wild type) equilibrium simulation (black curve) and the mutants K39E-M62E-Q66E-L68D-M43E (red) and M62E-Q66E-L68D (blue). A clear left-shift is observed as compared to the typical conformations observed for the intermediary and opened states, indicated by vertical lines. (*Bottom right*) The same distribution of opening angles is also shown for additional mutants simulated.
